# Toxicity and Biodegradation of Two Different Hydrothermal Liquefaction Process Waters to Anaerobic Digestion and the Effect of Microaeration

**DOI:** 10.1101/2024.12.10.627544

**Authors:** Mei Zhou, Joseph G. Usack, Aidan Mark Smith, Largus T. Angenent

## Abstract

Hydrothermal liquefaction (HTL) can convert a considerable portion of the carbon in complex feedstocks into renewable bio-oil, but it also generates a liquid byproduct (*i.e.*, HTL process water) that retains 15 – 55% of the carbon from the HTL feedstock. Feeding HTL process water to anaerobic digestion (AD) is a promising approach for maximizing resource recovery, enabling the conversion of the retained carbon into biogas. However, various toxic and poorly biodegradable compounds in HTL process water make its treatment with AD challenging. Presently, the underlying mechanisms remain often unclear. We investigated the impact of HTL process water from two different feedstocks – a food-waste proxy (*i.e.*, dog food, rich in proteins) and wheat straw (rich in lignocellulose) – on the different trophic groups in the food web of AD. We found that methanogens rather than acidogens were inhibited by HTL process water. Comparative toxicity and biodegradability analyses showed that wheat-straw process water had a higher biodegradability regardless of its higher toxicity to acetoclastic methanogens than food-waste process water, due to its higher content in toxic but easily degradable aromatic compounds. Microaeration enhanced the biodegradation and methane yields of food-waste process water, particularly under anoxic conditions. However, microaeration was ineffective for wheat-straw process water. These findings highlight the importance of feedstock-specific strategies to optimize AD for biogas production from HTL process water.

## 1 Introduction

Integrating hydrothermal liquefaction (HTL) and anaerobic digestion (AD) is a promising approach for maximizing resource recovery from wet biomass and organic waste [1–3]. HTL operates in hot compressed water, requiring an operating temperature of 250 – 374°C, a pressure of 5 – 20 MPa, and a reaction time of 5 – 60 min [4]. This process can convert complex feedstocks into bio-oil with a higher energy value at relatively high rates [5]. However, HTL process water is also produced, which is a liquid byproduct that retains 15 – 55% of the carbon from the HTL feedstocks [6]. Valorizing HTL process water through AD was shown to recover an additional 2 – 14% of the initial energy, thereby, enhancing the net energy gain of biomass conversion [7], while also utilizing a product that is a waste.

During AD, the carbon in HTL process water is converted into biogas (mainly methane and carbon dioxide) through three sequential trophic steps of acidogenesis, acetogenesis, and methanogenesis within the AD food web [8]. However, AD conversion faces considerable challenges due to the complex composition and potential inhibitory effects of HTL process water on the trophic microbial groups for AD. The feedstock for HTL treatment influences the composition and inhibition of the resultant process water. For instance, HTL process water from lignocellulosic feedstocks (*e.g.*, rice straw and corn stover) mainly comprises phenolic and furanic compounds, which have been reported to be toxic to AD microbes [9]. Reportedly, HTL process water from protein-rich feedstocks contains more N-heteroaromatic compounds, which are hard to degrade in strictly anaerobic conditions [10].

Other studies [11, 12] have indicated that the extent of inhibition directly correlates with the concentration of HTL process water and HTL operating temperatures, with higher concentrations and higher temperatures exacerbating the inhibitory effects. However, the inhibition mechanisms are still unclear. Further research is required to understand whether inhibition is due to high concentrations of hard-to-degrade components or toxic components in the HTL process water and whether any of the three trophic steps in the AD food web are particularly vulnerable to this inhibition. Understanding the underlying mechanisms and developing strategies to mitigate this inhibition is crucial to optimizing AD of HTL process water, thereby, enhancing not only energy resource recovery but also the degradation of environmentally harmful compounds.

One potential strategy to mitigate the inhibitory effects and improve the biodegradability of HTL process water is microaeration within the anaerobic digester. Introducing small amounts of oxygen or air during AD has proven to increase microbial diversity, leading to more efficient substrate degradation and methane production [13]. Oxygen addition promotes the growth of facultative anaerobes, which can function in both aerobic and anaerobic conditions, facilitating the breakdown of complex organic compounds into simpler molecules that are more accessible to strictly anaerobic microbes [14]. Additionally, microaeration enables the limited aerobic respiration to generate extra ATP, which boosts microbial metabolism and accelerates substrate degradation [15]. Although microaeration has been effective in other applications, it has not been specifically tested for HTL process-water treatment.

This study aims to elucidate the impact of feedstock composition and microaeration on the toxicity and biodegradability of HTL process water during AD. By examining HTL process water from two dissimilar feedstocks – a food-waste proxy (*i.e.*, dog food, rich in proteins) and wheat straw (rich in lignocellulose) – this research evaluates how different chemical compositions of HTL process water affect key microbial processes in AD, particularly methanogenesis. Additionally, the study explores the potential of microaeration-acclimated biomass as a strategy to tolerate toxicity and enhance the biodegradability of HTL process water. This research provides valuable insights into optimizing AD processes for treating HTL process water and improving its environmental sustainability.

## 2 Materials and methods

### 2.1 Characteristics of HTL process water and inoculum

We generated two HTL-process-water solutions: food-waste process water (FWPW) and wheat-straw process water (WSPW) with the Aarhus University pilot-scale continuous HTL reactor at AU Viborg (Denmark) (**Table S1**). Dog food was selected as a proxy feedstock for protein-rich food wastes (Orlando Multico, Lidl, Denmark), while wheat straw was a model feedstock for lignocellulose-rich biomass. The HTL process conditions were 350 for 15 min with 12% dry matter. After gravimetric bio-oil separation, ten 10-L bottles of each process-water solution were shipped to our laboratory at the University of Tübingen in Germany. There, each process-water solution was filtered using 5-μm filter bags, stirred, combined in a large container, and redistributed again into 10-L bottles. The HTL-process-water solutions were stored at -20 until further use.

For anaerobic toxicity (AT) assays, we collected the anaerobic inoculum from an anaerobic digester that was treating animal manure at Hohenheim University (Reutlingen, Germany). For the acetoclastic methanogenic activity (AMA) assays and the biochemical methane potential (BMP) assays, we utilized biomass from an active 5.5-L laboratory AD reactor (fully anaerobic) and a microaeration-acclimated AD reactor to study the toxicity and biodegradability of HTL process water. The only operating difference between the two laboratory reactors was intermittent air dosing at 15.6 mL/min into the microaeration reactor. The biomass had acclimated to microaeration with air for one year. All assays were performed in a near-neutral pH range (6.9 – 7.3). The sufficient alkalinity of the inoculum made pH adjustments unnecessary.

### 2.2 Anaerobic toxicity (AT) assays

The classical AT assays to measure toxicity involved introducing varying chemical oxygen demand (COD) loads of HTL process water to the serum bottles containing a consistent amount of standard substrate and anaerobic inoculum [16, 17]. All assays were performed in triplicate at 37°C in 250-mL glass serum bottles with a liquid volume of 100 mL. We also performed control assays without HTL-process-water interference. The COD load of HTL process water was expressed as a percentage of the standard substrate COD. For instance, the FWPW_10% scenario (**Table S2, S3**) indicated that the COD added by the FWPW accounted for 10% of the standard substrate COD. WSPW_10% (**Table S2, S3**) indicated that the COD added by the WSPW accounted for 10% of the standard substrate COD. For the acidogenic AT assay (**Table S2**), we used a glucose solution (1 g/L) as the standard substrate to examine the impact of HTL process water on the acidogenesis in the AD food web [18]. We assumed that a reduction in glucose degradation rates relative to the control scenario indicated acidogenesis inhibition. We measured the glucose, lactate, short-chain carboxylic acid (SCCA), and medium-chain carboxylic acid (MCCA) concentrations during incubation at 0, 6.5, 10.5, 22, 31, 49.5, 81.5, 105, and 127 h. For the methanogenic AT assay (**Table S3**), we followed the method described by Owen *et al.*[16], which used a mixture of acetate and propionate as the standard substrate. We assumed that a reduction in methane production rates relative to the control scenario indicated methanogenesis inhibition. Biogas production and composition were measured daily to monitor the process.

### 2.3 Acetoclastic methanogenic activity (AMA) assays

Because acetoclastic methanogens are recognized as the dominant methanogenic group (*i.e.*, 80 – 95% relative abundance) compared to hydrogenotrophic methanogens during AD [19, 20], we performed (AMA) assays in 60-mL serum bottles to measure the toxicity of different reactants. Reactants included HTL process water from two dissimilar feedstocks (as complex reactants) and specific inhibitory compounds (as defined reactants) typically found in HTL process water. Each 60-mL serum bottle for the AMA assay contained 30 mL of liquid, consisting of anaerobic water, 15.8 mL sodium-acetate solution (2 g/L) as standard substrate, 2.8 mL biomass, and reactants, along with 0.3 mL of trace-element solution, mineral-stock solution, and Na_2_S solution (10 g/L). The trace-element solution consisted of (in g/L): C_6_H_9_NO_6_, 2; MnSO_4_, 1; FeSO_4_, 0.8; CoCl_2_, 0.2; ZnSO_4_, 0.2; CuCl_2_, 0.02; NiCl_2_, 0.02; Na_2_MoO_4_*2H_2_O, 0.02; Na_2_SeO_4_, 0.02; and Na_2_WO_4_, 0.02. The mineral stock contained the following (in g/L): NaCl_2_, 40; NH_4_Cl, 50; KCl, 5; KH_2_PO4, 6; MgCl_2_, 7; and CaCl_2_, 2.

After inoculation, the bottles were sparged with nitrogen gas for 10 min. Subsequently, the bottles were placed in a 37 shaking incubator (150 rpm) for 20 h. To mitigate the initial lag phase, 8 mL of sodium acetate (2 g/L) solution was supplemented to each bottle at 21 h [21]. Bottles were resealed, sparged with nitrogen gas, and returned to the incubator. The AMA was calculated and expressed as g COD-CH_4_/g VSS/day according to Equations (**Eq.** 1 – 3):

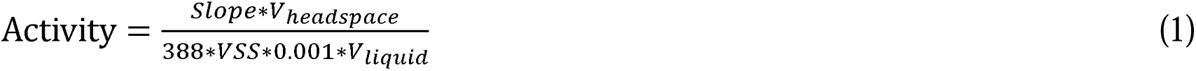

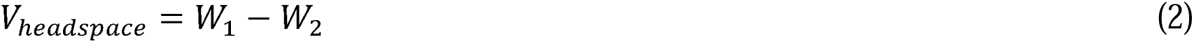

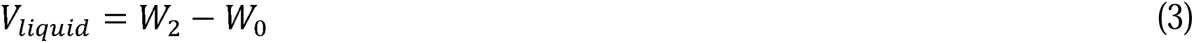

where *1/388* is the COD value of 1 mL CH_4_ at 25□; *Slope* is the slope of the curve for methane composition in the headspace versus time (% CH_4_/day); *VSS* is the volatile suspended solids at the end of the tests (g VSS/L); *V_headspace_* and *V_liquid_* are the headspace gas volume and the liquid volume, respectively (mL); *W_0_*, *W_1_*, and *W_2_*are the weights of the blank bottle, the bottle full of distilled water, and the bottle with samples in the end, respectively (g). The density of the liquid was assumed to be 1 g/cm^3^.

### 2.4 Biochemical methane potential (BMP) assays

We conducted BMP assays to determine the biodegradability of HTL process water from different feedstocks. Every BMP assay was performed in triplicate in 60-mL septum-capped bottles with a total liquid volume of 30 mL, consisting of HTL process water at a target COD of 1 g/L and anaerobic biomass with an inoculum/substrate (I/S) ratio of 3.5 (on a COD basis). Additionally, we added the same amount of trace-element solution, mineral-stock solution, and Na_2_S solution (10 g/L), as used in the AMA assays in Section 2.3. After inoculation, the bottles were flushed with nitrogen gas and incubated at 37□ in a shaking incubator (150 rpm). To test the efficacy of microaeration as a biodegradation strategy at two dosing intensities, we replaced nitrogen in the headspace with air to achieve oxygen partial pressures of 1% and 3% (*v/v*) (**Table S4**).

Biogas production was determined using the syringe method, which involved measuring the biogas volume with a glass syringe inserted directly through the septum of the bottle lid. The overpressure inside the incubated bottles pushed the piston until it balanced with atmospheric pressure. The biogas was then released into a fume hood. The measured biogas volume was normalized to standard conditions (*i.e.*, 25□, 101.3 kPa). The biogas volumes measured in the blank bottles were substrated from the biogas volume measured in the experimental bottles. The cumulative methane production curves were simulated by the modified Gompertz equation [22], as presented in Equation (**Eq. 4**).

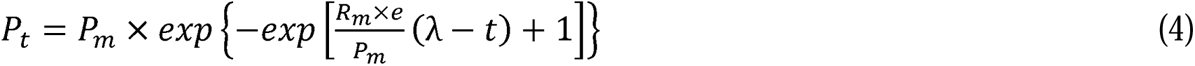

where *P_t_* is the methane production volume at time t (mL); *P_m_* is the maximum biomethane yield potential (mL); *R_m_* is the peak methane production rate (mL/day); λ is the duration of the lag phase (day); *t* is the incubation period (day); *e* is the mathematical constant approximately equivalent to 2.7. Origin 8.5 software was used to perform a nonlinear regression analysis to determine the parameters of *P_m_*, *R_m_*, λ, and the correlation coefficient of determination (R^2^).

### 2.5 Analytical methods and data analysis

The COD was analyzed according to Standard Methods [23]. SCCAs and MCCAs were determined using a gas chromatograph (GC), which was equipped with a thermal conductivity detector (TCD) (7890B GC System, Agilent, USA) using a capillary column (Nukol, 15m X 0.25 mm I.D. X 0.25 μm). Glucose and lactate were determined using a high-performance liquid chromatograph (HPLC) (Shimadzu LC 20AD), which was coupled with a refractive index and UV detector (Shimadzu, Kyoto, Japan). Biogas composition was measured using a GC (SRI gas GCs, SRI Instruments, USA) equipped with a 0.3-m HaySep-D packed Teflon column and a flame ionization detector using an isothermal oven temperature of 70°C. Pure methane was used as the standard for GC, and gas samples of AMA assays were analyzed hourly to plot the percentage of methane in the headspace during the incubation period.

### 2.6 Statistical Analysis

Data from duplicate/triplicate measurements were expressed as mean ± standard error. All statistical analyses were performed using the OriginPro student version (OriginLab, Northampton, USA). Values of p < 0.05 were considered to be statistically significant.

## 3 Results and discussion

### 3.1 Methanogenesis, rather than acidogenesis, is more severely inhibited during AD of HTL process water, as shown with anaerobic toxicity (AT) assays

To determine which trophic step in the AD food web is most susceptible to HTL-process-water toxicity, we monitored glucose conversion during acidogenesis and methane production during methanogenesis for AT assays. We did not measure the activity of acetogenesis because we could not isolate this microbial activity due to overlapping activity between acidogenesis and methanogenesis [24]. The introduction of HTL process water did not markedly inhibit acidogenesis. Glucose was fully degraded within 22 h in all assays regardless of the HTL process water COD load (**Fig. 1a, 1b**). Lactate was the primary intermediate produced from glucose utilization. SCCAs and small amounts of MCCAs were also produced, and their concentration increased during the incubation period for each scenario (FWPW_0% – WSPW_100% in **Fig. 1c**, **1d**). Lactate concentrations decreased after ∼ 31 h for all scenarios (**Fig. 1a**, **1b**), while acetate and propionate concentrations increased (**Fig. 1c**, **1d**), suggesting lactate oxidation had occurred.

**Fig. 1.**
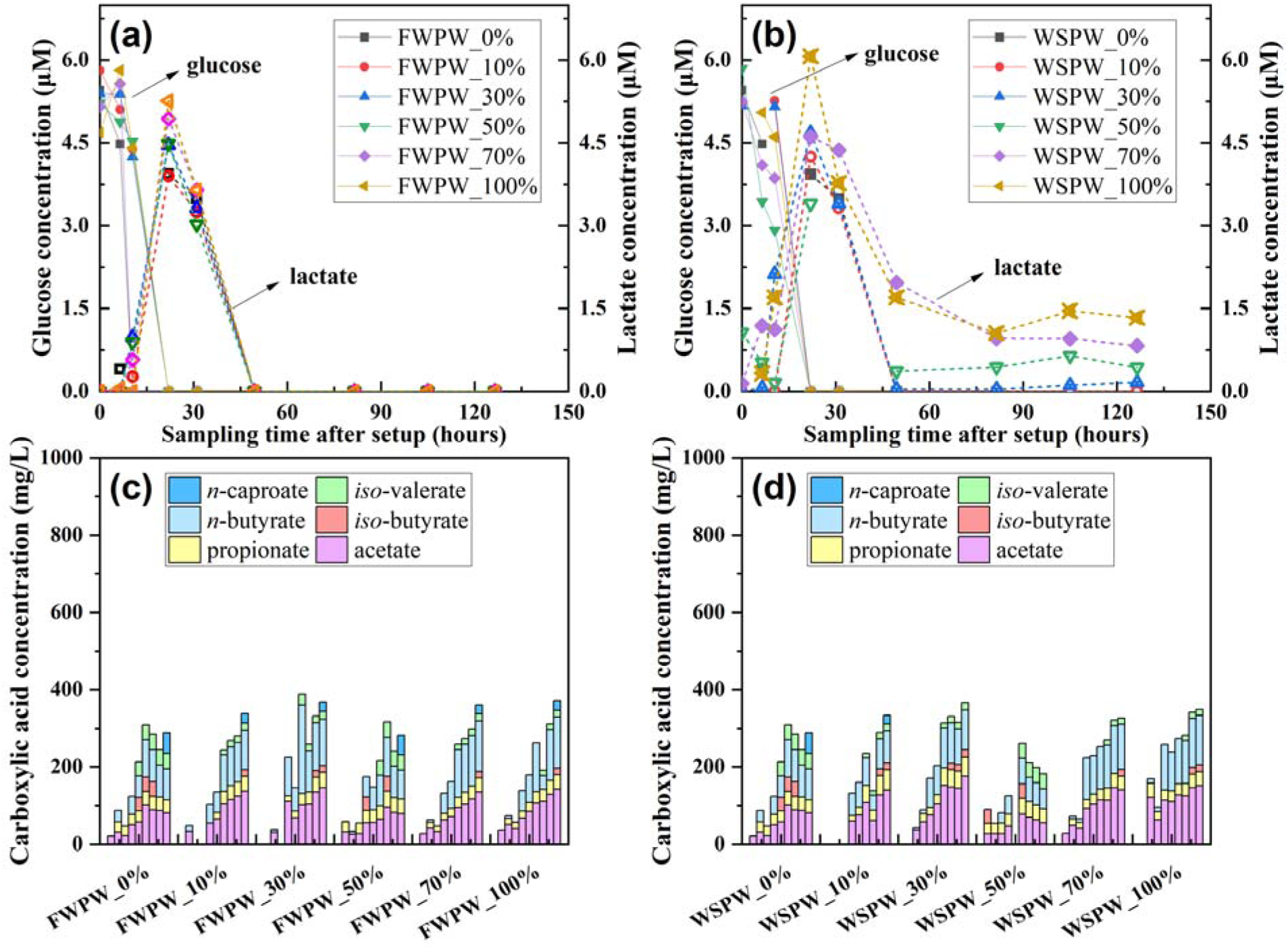
Anaerobic toxicity (AT) assays for effects of food-waste process water (FWPW) and wheat-straw process water (WSPW) on acidogenesis. Glucose and lactate concentrations (**a, b**) and carboxylic acid concentrations (**c, d**). The increasing levels of HTL process water added as a feed are expressed as percentages of the standard substrate COD (*i.e.*, glucose). For each condition, sampling times were 0, 6.5, 10.5, 22, 31, 49.5, 81.5, 105, 127 h after setup. We analyzed the carboxylic acid concentration of scenarios of FWPW_0%, FWPW_50%, WSPW_0%, and WSPW_0% in the same running sequence and analyzed scenarios of FWPW_10%, FWPW_30%, FWPW_70%, FWPW_100%, WSPW_10%, WSPW_30%, WSPW_70%, and WSPW_100% in the other running sequence.

Lactate oxidation into acetate, propionate, and H_2_ precedes methanogenesis [25, 26]. The H_2_ produced from lactate and propionate oxidation must be removed by hydrogenotrophic methanogens and maintained at low partial pressures for the oxidation reaction to occur. Lactate was completely oxidized in all scenarios involving food-waste process water (**Fig. 1a**). However, lactate oxidation was incomplete at higher COD loads of wheat-straw process water (*i.e.*, > WSPW_30%) (**Fig. 1b**), potentially due to the inhibition of lactate-utilizing bacteria, acetogens, or hydrogenotrophic methanogens. The decreased acetate accumulation rate for the scenarios > WSPW_30% (decreased slope of pink bars in **Fig. 1d**) compared to scenarios < WSPW_50% also indicated the inhibition of lactate conversion by higher concentrations of wheat-straw process water (**Fig. 1d**).

Digestion with HTL process water from different feedstocks resulted in similar inhibition trends during methanogenesis (**Fig. 2)**. According to the time course of cumulative methane production, longer lag phases occurred at higher COD loads of HTL process water, especially in the FWPW_50%, FWPW_70%, FWPW_100%, WSPW_70%, and WSPW_100% scenarios (**Fig. 2a**, **2b**). Methane production gradually increased after 6 days of incubation, possibly due to biomass acclimatization or the degradation of inhibitory compounds [9]. The total cumulative methane production was greater for higher COD loads because more COD was added and degraded.

**Fig. 2.**
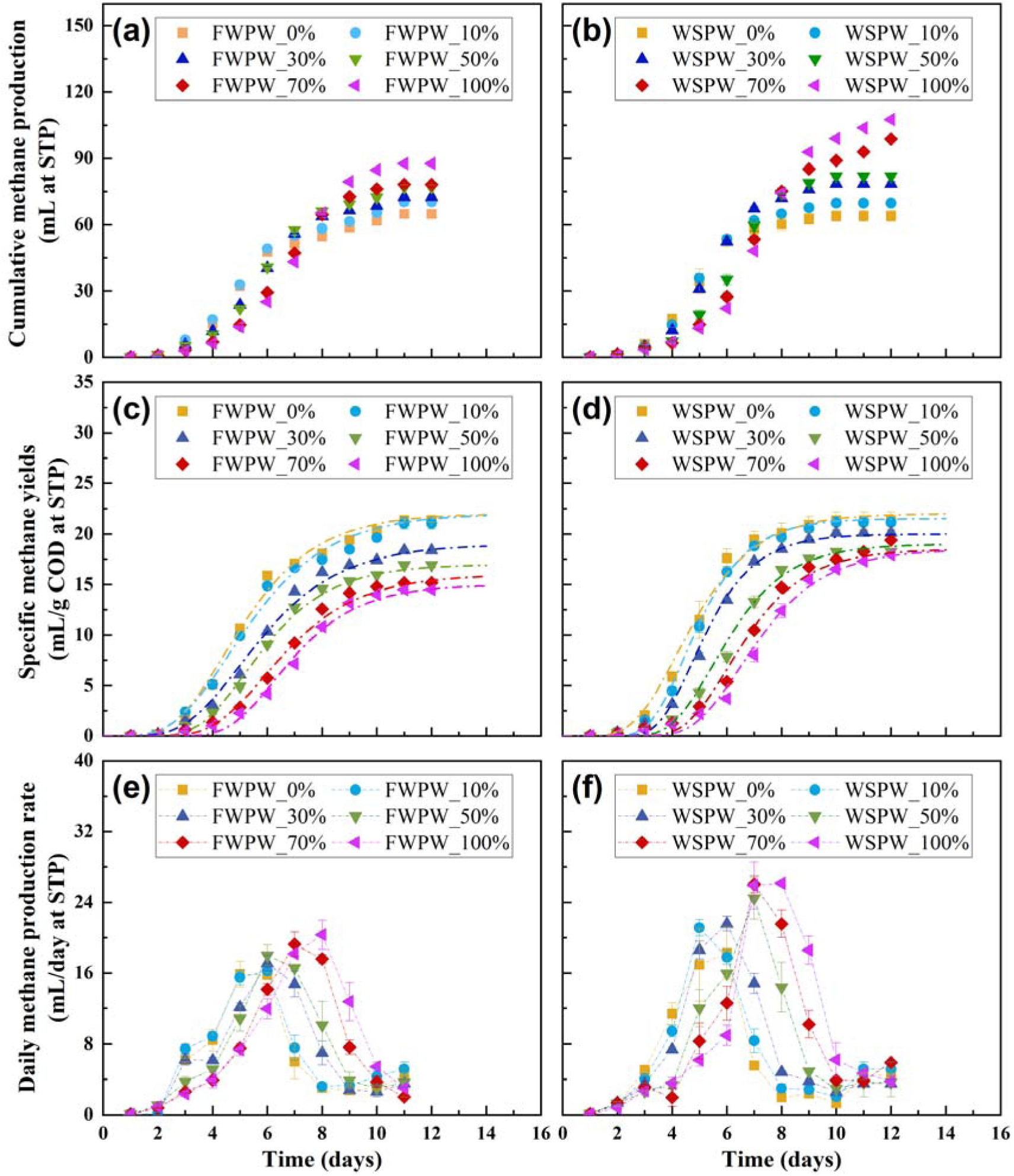
Anaerobic toxicity (AT) assays for effects of food-waste process water (FWPW) and wheat-straw process water (WSPW) on methanogenesis: Cumulative methane yields (mL at STP) (**a, b**), specific methane yields (mL/g COD at STP) (**c, d**), and daily methane production rate (mL/day) (**e, f**). The increasing levels of HTL process water added as a feed are expressed as percentages of the standard substrate COD (*i.e.*, the mixture of sodium acetate (10 g/L) and propionic acid (6.69 mL/L)). Data for each experimental condition are based on replicate experiments using separate bottles. Statistical analysis was performed using Two-Way ANOVA, and the results are presented as mean ± standard deviation (data points + error bars). Some error bars may not be fully visible due to overlapping data points; detailed numerical ranges can be found in Table S5.

Specific methane yields decreased at higher COD loads of HTL process water as anticipated (**Fig. 2c**, **2d**). The assays with FWPW_10% and WSPW_10% achieved the highest specific methane yields with HTL process exposure, achieving 21.1 ± 0.57 and 20.8 ± 0.11 mL STP CH_4_/g COD, respectively. However, when compared to the control assays (only acetate and propionate substrate), no differences in methane yield were detected (p > 0.05), indicating no apparent inhibition of methanogenesis at HTL-process-water COD loads at 10%. When COD loads exceeded 30%, methane yields were lower than the controls (**Fig. 2c**, **2d**), suggesting that the process water was inhibitory or less biodegradable than the control substrate. The daily methane production rates showed earlier peak values upon adding FWPW_10% and WSPW_10%, while the peak values were delayed but increased at higher COD ratios of HTL process water (**Fig. 2e**, **2f**).

To better compare the toxicity of HTL process water from different feedstocks, the time courses of specific methane yield curves (in **Fig. 2c**, **2d**) were modeled using the modified Gompertz equation (**Eq. 4**). The simulated results aligned well with the measured impacts of higher COD loads of HTL process water on maximum methane yields (P_m_), maximum daily methane production rate (R_m_), and the lag phase (λ) (**Table 1**). According to the model, food-waste process water had a more adverse effect on the maximum methane yields and maximum daily methane production rates when compared with wheat-straw process water (**Table 1**). Additionally, the lag phases for food-waste process water and wheat-straw process water were up to two days longer than the control (**Table 1**).

**Table 1.** Experimental and kinetic model parameters simulated by the modified Gompertz equation for food-waste process water (FWPW) and wheat-straw process water (WSPW).

| Substrates | Measured maximum | Kinetic model parameters |  |  |
| --- | --- | --- | --- | --- |
| | methane yields<br>(mLCH<sub>4</sub>/gCOD<sub>added</sub>) | $P_m$<br>(mLCH<sub>4</sub>/gCOD<sub>added</sub>) | $R_m$<br>(mLCH<sub>4</sub>/gCOD<sub>added</sub>/day) | $\lambda$<br>(day) |
| FWPW_0% | 21.4 ± 0.20 | 22 | 12 | 2.6 |
| FWPW_10% | 21.1 ± 0.57 | 22 | 11 | 2.6 |
| FWPW_30% | 18.4 ± 0.08 | 19 | 9.5 | 3 |
| FWPW_50% | 16.9 ± 0.20 | 17 | 10 | 3.5 |
| FWPW_70% | 15.2 ± 0.19 | 16 | 8.5 | 4 |
| FWPW_100% | 14.5 ± 0.04 | 15 | 9 | 4.6 |
| WSPW_0% | 22.6 ± 0.60 | 22 | 14 | 2.6 |
| WSPW_10% | 20.8 ± 0.11 | 21.5 | 17 | 3.2 |
| WSPW_30% | 19.8 ± 0.04 | 20 | 16 | 3.5 |
| WSPW_50% | 18.1 ± 0.07 | 19 | 12.8 | 4 |
| WSPW_70% | 18.1 ± 0.02 | 18.5 | 12 | 4.5 |
| WSPW_100% | 17.1 ± 0.01 | 18.5 | 11 | 4.8 |

P<sub>m</sub>: simulated maximum methane yields; R<sub>m</sub>: simulated maximum daily methane production rate; λ: lag phase.

The results from the acidogenesis and methanogenesis AT assays suggest that HTL process water more severely inhibits methanogenesis than acidogenesis (**Fig. 1**, **2**). These results agree with findings from another study [27], which reported reduced methane production when HTL process-water concentration from algal biomass exceeded 40% (*v/v*) in the feed. Interestingly, Macêdo *et al*. [28] observed an increased relative abundance of methanogenic archaea but still noted SCCA accumulation and cessation of methane production at the highest concentration of HTL process water from primary sewage sludge.

### 3.2 HTL process water type and concentration differentially hinder acetoclastic methanogenic activity, as shown with anaerobic methanogenic activity (AMA) assays

From the analysis of AT assay results, both HTL process water types inhibited methane production when the COD load was equal to or higher than 30% of the standard substrate (**Fig. 2**). To understand the inhibition mechanisms better, we also conducted short-term (2-day) AMA assays to assess the toxicity to acetoclastic methanogens from different concentrations of food-waste process water and wheat-straw process water. In addition, we also assessed the toxicity to acetoclastic methanogens from specific inhibitory compounds, which are common in process water. The second acetate dosing in each assay, which was administered after approximately one day of incubation, primarily contributed to methane production due to its fast kinetics (and thus not the added process water or compounds) [29]. The initial incubation period of one day allowed for the adaptation and stabilization of biomass in response to acetate as a substrate. The pH level for all bottles was within the neutral range of 6.9 – 7.3 throughout the two-day period, thanks to the high alkalinity of the inoculum.

Food-waste process water and wheat-straw process water influenced acetoclastic methanogenic activity differently at elevated concentrations (p < 0.01 in **Fig. 3a**). Food-waste process water did not significantly reduce acetoclastic methanogenic activity when the COD load was up to 2 g/L (p > 0.05 in **Fig. 3a).** Notably, the toxicity of wheat-straw process water was more severe with the increased concentrations (p < 0.01 in **Fig. 3a**). The half maximal inhibitory concentrations (IC_50_), which corresponds to 50% inhibition, was 2.5 g COD/L for wheat-straw process water, but considerably higher for food-waste process water and outside of our tested concentrations (> 5 g COD/L in **Table 2**). This difference in toxicity response can be attributed to compositional differences between the two HTL process water types. Nitrogen-containing compounds, including pyridines, pyrazines, pyrimidines, and pyrroles, dominate HTL process water from protein-rich feedstocks [30]. In contrast, lignocellulosic feedstocks generate HTL process water rich in oxygen-containing compounds, including phenols, ketones, and furans [31].

**Fig. 3.**
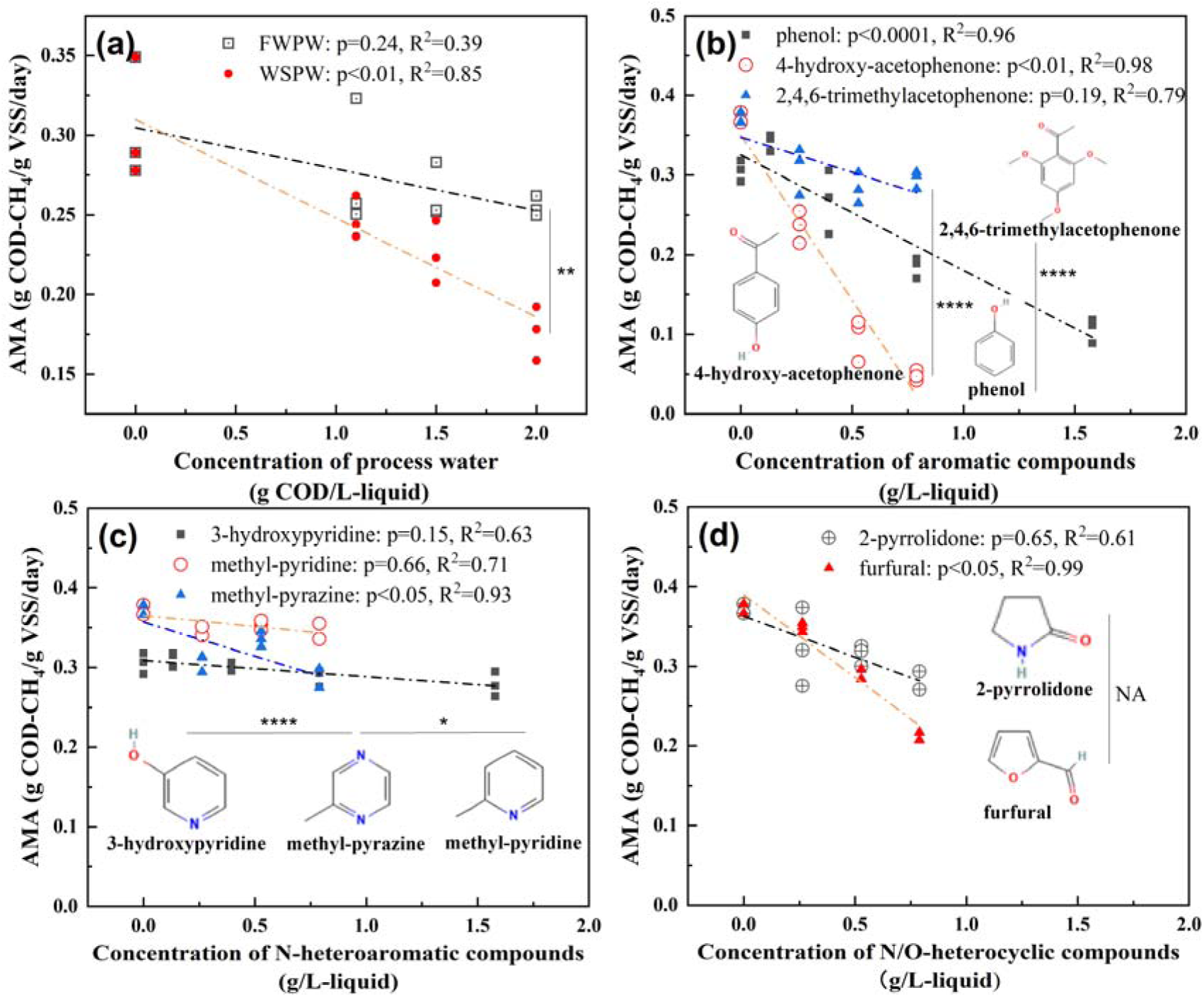
Acetoclastic methanogenic activity (AMA) assays for the effect of different reactants on acetoclastic methanogenic activity (expressed as g COD-CH_4_/g VSS/day). Reactants include the increasing concentrations of food-waste process water (FWPW) and wheat-straw process water (WSPW) (**a**) and specific inhibitory compounds, including aromatic compounds (**b**), N-heteroaromatic compounds (**c**), and N/O-heterocyclic compounds (**d**). We conducted One-Way ANOVA for each reactant (p-value in the legend) to assess the effect of elevating concentrations on AMA. We also assessed the significance between AMA and different reactants across concentrations by using Two-Way ANOVA (p-value with asterisks in the figure, * indicates p < 0.05, **** indicates < 0.0001, and NA is no significant difference).

**Table 2.** The 50% inhibition concentration (IC_50_) of process water and typical organic inhibitors on acetoclastic methanogenic activity as shown with the AMA assay.

| Inhibitors | 50% Inhibition Concentration (IC<sub>50</sub>) |
| --- | --- |
| Food-waste process water | Outside our range, likely > 5 g COD/L |
| Wheat-straw process water | 2.5 g COD/L |
| 4-hydroxy-acetophenone | 0.42 g/L |
| Furfural | 0.94 g/L |
| Phenol | 1.1 g/L |
| 2,4,6-trimethoxyacetophenone | 2.0 g/L |
| 2-pyrrolidinone | 1.8 g/L |
| Methyl-pyrazine | Outside our range, likely > 4 g COD/L |
| Methyl-pyridine | Outside our range, likely > 6 g COD/L |
| 3-hydroxypyridine | Outside our range, likely > 7 g COD/L |

Some studies reported that high concentrations of nitrogen-containing compounds in HTL process water may inhibit methane production [25, 27]. However, this indication is speculative because the studies relied on the cytotoxicity data by Pham *et al.* [32]. These authors demonstrated that pyridine and pyrrolidine compounds have high lethal concentration 50 (LC_50_) values, reducing the cell density of Chinese hamster ovary cells by 50% at concentrations of 5500 µM and 16,900 µM, respectively. It is important to note that Chinese hamster ovary cells are very different from the microbes involved in the AD process. In addition to nitrogen-containing compounds, other studies have suggested that aromatic compounds (*e.g.*, phenols and furans) may inhibit acetogenesis and methane production during AD of HTL process water from swine manure [5, 33]. Still, the effects of these potential inhibitors are not yet well understood.

The molecular structure of inhibitors can be an important factor in inhibiting acetoclastic methanogenic activity to varying degrees, and therefore we measured this activity (**Fig. 3b, 3c, 3d**). Here, we found, at the same concentration ranges, that aromatic compounds (no N or O in the ring structure) with a hydroxyl group (4-hydroxy-acetophenone and phenol) showed a higher toxicity to acetoclastic methanogenic activity than 2,4,6-trimethoxyacetophenone without a hydroxyl group (p < 0.0001 in **Fig. 3b**). For N-heteroaromatic compounds, the number of heteroatoms correlated with the degree of toxicity to acetoclastic methanogens. For instance, methyl-pyrazine (2N in the ring) reduced the acetoclastic methanogenic activity more severely than methyl-pyridine (p < 0.05) and 3-hydroxypyridine (1N in the ring) (p < 0.0001 in **Fig. 3c**).

Additionally, for the non-aromatic heterocyclic compounds, the toxicity of furfural (with O as the heteroatom) was significantly influenced by the concentration (p < 0.05), whereas the toxicity of pyrrolidone (with N as the heteroatom) was not significantly affected by the concentration (p > 0.05 in **Fig. 3d**). In general, higher toxicity to acetoclastic methanogens was found for the individual compounds that can be anticipated to be more abundant in wheat-straw process water, such as the aromatic compounds and furfural without N in their aromatic rings, compared to food-waste process water, such as the N-heteroaromatic compounds and 2-pyrrolidone with N (**Fig. 3b, 3c, 3d** and **Table 2**). This finding is consistent with the higher toxicity that we observed for wheat-straw process water than food-waste process water.

### 3.3 HTL process water from protein-rich feedstock is less biodegradable than that from lignocellulose-rich feedstock, as shown with biological methane potential (BMP) assays

For the BMP assays, HTL process water was added to the bottles at a COD concentration of 1 g/L to measure biodegradability at a very mild toxicity level. Indeed, AMA assays showed that adding 1 g COD/L of food-waste process water and wheat-straw process water resulted in only 8.5% and 20% reduction in acetoclastic methanogenic activity, respectively (**Fig. 3a**). Other indications of relatively low toxicity levels for HTL process water at a concentration of 1 g COD/L are: **(1)** the methane production increased rapidly in the initial phase without a lag phase (**Fig. 4a**); and **(2)** all SCCAs remained below the detection level (∼ 0.47 mM) during the BMP-assay period of 30 days.

**Fig. 4.**
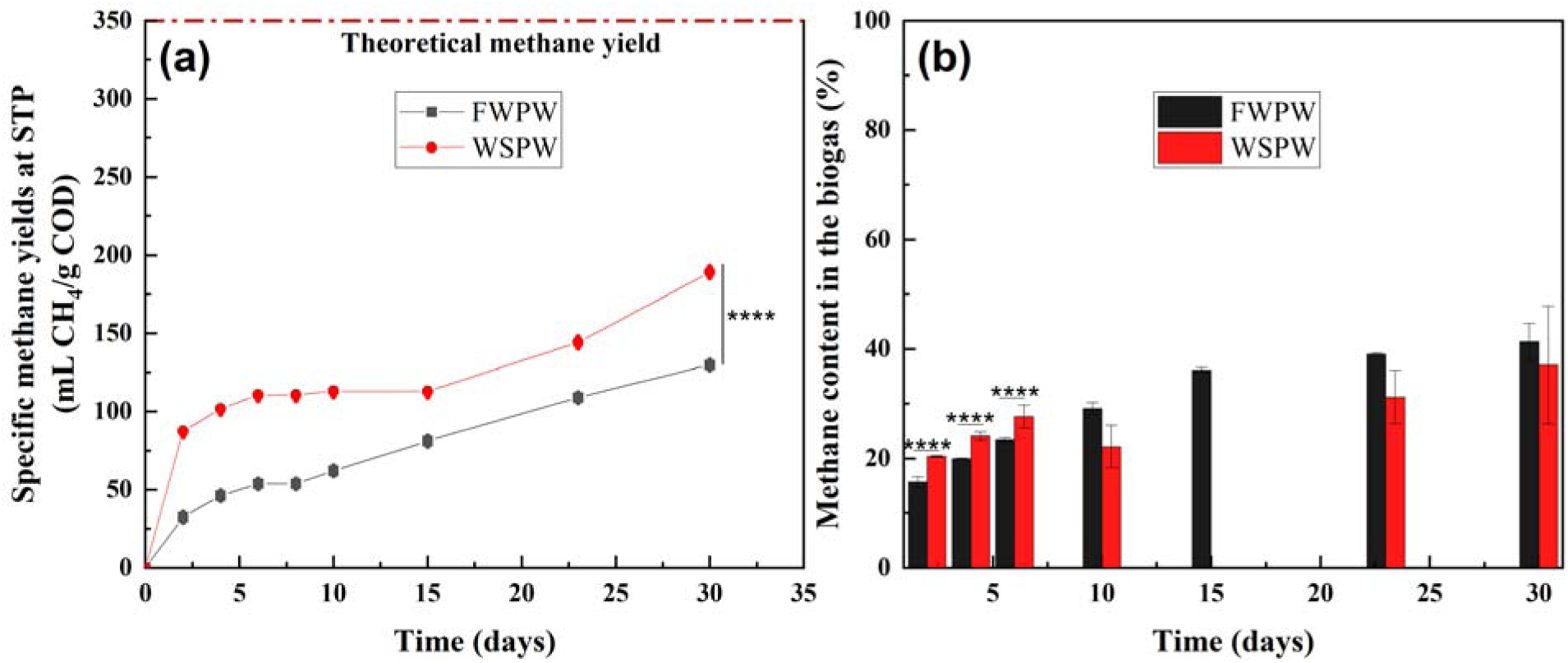
Biochemical methane potential (BMP) assays for specific methane yields of HTL process water (**a**) and methane content in the biogas (**b**) from food-waste process water (FWPW) and wheat-straw process water (WSPW). The methane production from the anaerobic inoculum was subtracted to calculate the specific methane yields from the two process water types. Data for each experimental condition are based on duplicate experiments using separate bottles. The results are presented as mean ± standard error (data points + error bars) and compared with the Paired-Sample t-Test (p-value with asterisks in the figure, **** indicates < 0.0001). Some error bars may not be fully visible due to overlapping data points; detailed numerical ranges can be found in Table S6.

The biodegradabilities of food-waste process water and wheat-straw process water were considerably different, as indicated by their respective specific methane yields (**Fig. 4a**). After 30 days of anaerobic digestion, the specific methane yields were 130 ± 0.13 and 1.9×10^2^ ± 1.5 mL CH_4_/g COD for food-waste process water and wheat-straw process water, respectively (p < 0.0001 in **Fig. 4a**). A lower biodegradability of food-waste process water than wheat-straw process water was also shown by a lower methane content in the biogas during the initial phase when the methane content was raising and not at steady state yet (p < 0.0001 in **Fig. 4b**). The lower biodegradability of food-waste process water could be due to the presence of recalcitrant N-heteroaromatic compounds, such as pyridines and pyrazines, which are produced from protein during HTL [34, 35].

Notably, wheat-straw process water had a higher methane potential than food-waste process water during the whole digestion period despite it showing more significant toxicity to acetoclastic methanogens, as discussed in Section 3.2. In two different publications after the same HTL treatment at 320°C for 0.5 h, Chen *et al.* also reported that HTL process water from rice straw (similar to our lignocellulose-rich wheat straw) had a higher biological methane potential of 217 mL CH_4_/g COD) [31] compared to that from protein-rich dewatered sewage sludge (similar to our protein-rich food-waste proxy), which yielded 136 mL CH_4_/g COD [36]. Given our toxicity and biodegradability results, improving process water biodegradability would be a more practical approach to enhance AD treatment than removing toxic compounds, such as through adsorption, especially for HTL process water from protein-rich feedstocks.

### 3.4 Microaeration-acclimated biomass enhances the biodegradability of one HTL process water type, as shown with the biological methane potential (BMP) assays

We tested the tolerance of microaeration-acclimated biomass to the two HTL process water types with AMA assays without applying any oxygen. Food-waste process water at concentrations lower than 2 g COD/L did not show a notable toxicity difference between microaeration-acclimated biomass and anaerobic biomass (p > 0.05 in **Fig. 5a**). When exposed to wheat-straw process water, microaeration-acclimated biomass exhibited a lower acetoclastic methanogenic activity compared with the anaerobic biomass (p > 0.0001 in **Fig. 5b**), showing a slight reduction of acetoclastic methanogen tolerance to wheat-straw process water with *prior* microaeration acclimation.

**Fig. 5.**
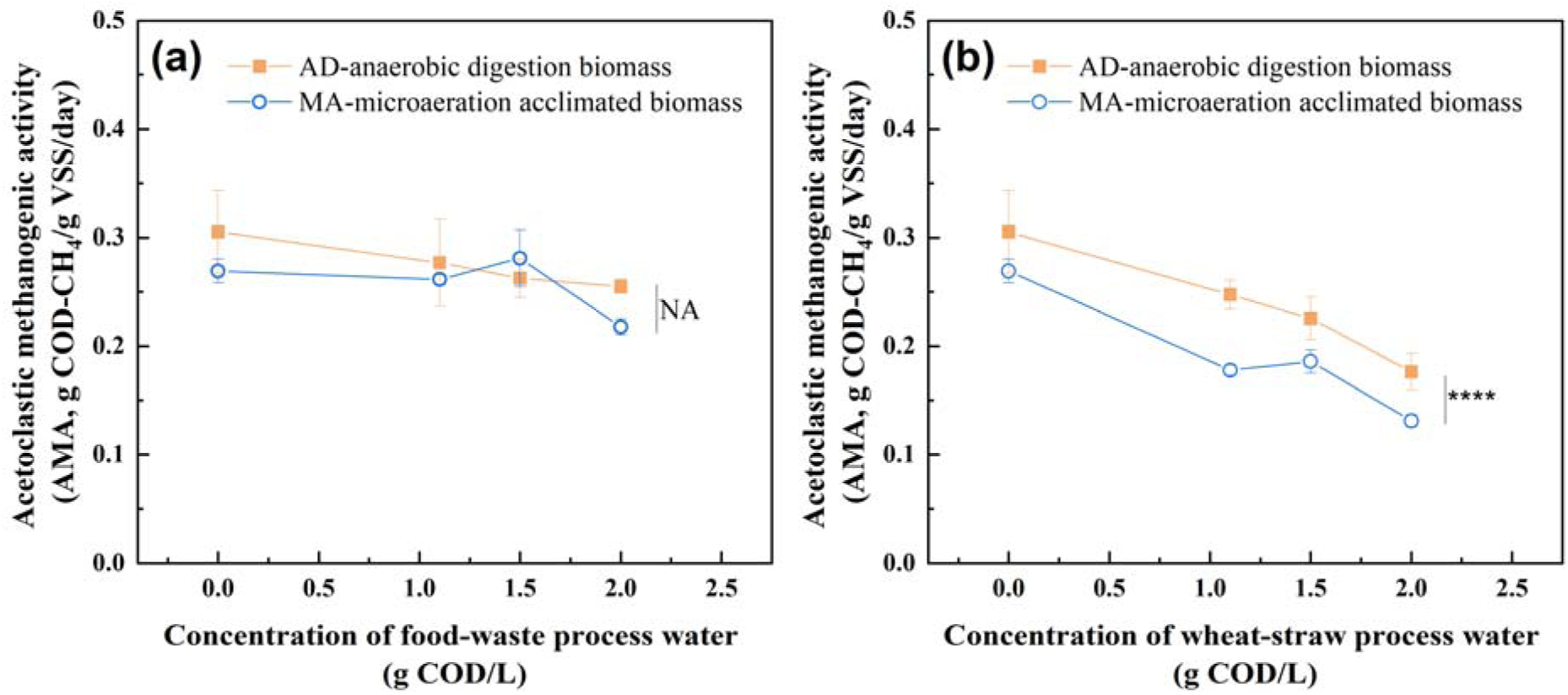
Acetoclastic methanogenic activity (AMA) assays for the effect on the activity of anaerobic digestion (AD) and microaeration-acclimated (MA) biomass with different concentrations of food-waste process water (FWPW) (**a**) and wheat-straw process water (WSPW) (**b**). All AMA bottles were kept strictly anaerobic. Data for each experimental condition are based on triplicate experiments using separate bottles. Statistical analysis was performed using Two-Way ANOVA, and the results are presented as mean ± standard deviation (data points + error bars) (p-value with asterisks in the figure, **** indicates < 0.0001, NA indicates no significant difference).

In addition to toxicity tolerance, we tested the ability of microaeration-acclimated biomass to biodegrade the two HTL process water types while varying the oxygen partial pressure in the headspace (**Fig. 6**). It is evident that microaeration-acclimated biomass consistently produced more methane from food-waste process water compared to anaerobic biomass (p < 0.0001 in **Fig. 6a, 6c, 6e**), even without the addition of oxygen (**Fig. 6a**). For wheat-straw process water, however, the specific methane yields were not significantly different (p > 0.05 in **Fig 6b**, **6d**, **6f**). We also observed for both HTL process water types that increasing the O_2_ partial pressures lowered the specific methane yields for each biomass, even for the microaeration-acclimated biomass (**Fig. 6**). Thus, for the food-process water for which the biodegradability was lower than wheat-straw process water, microaeration-acclimated biomass only had a biodegradability effect.

**Fig. 6.**
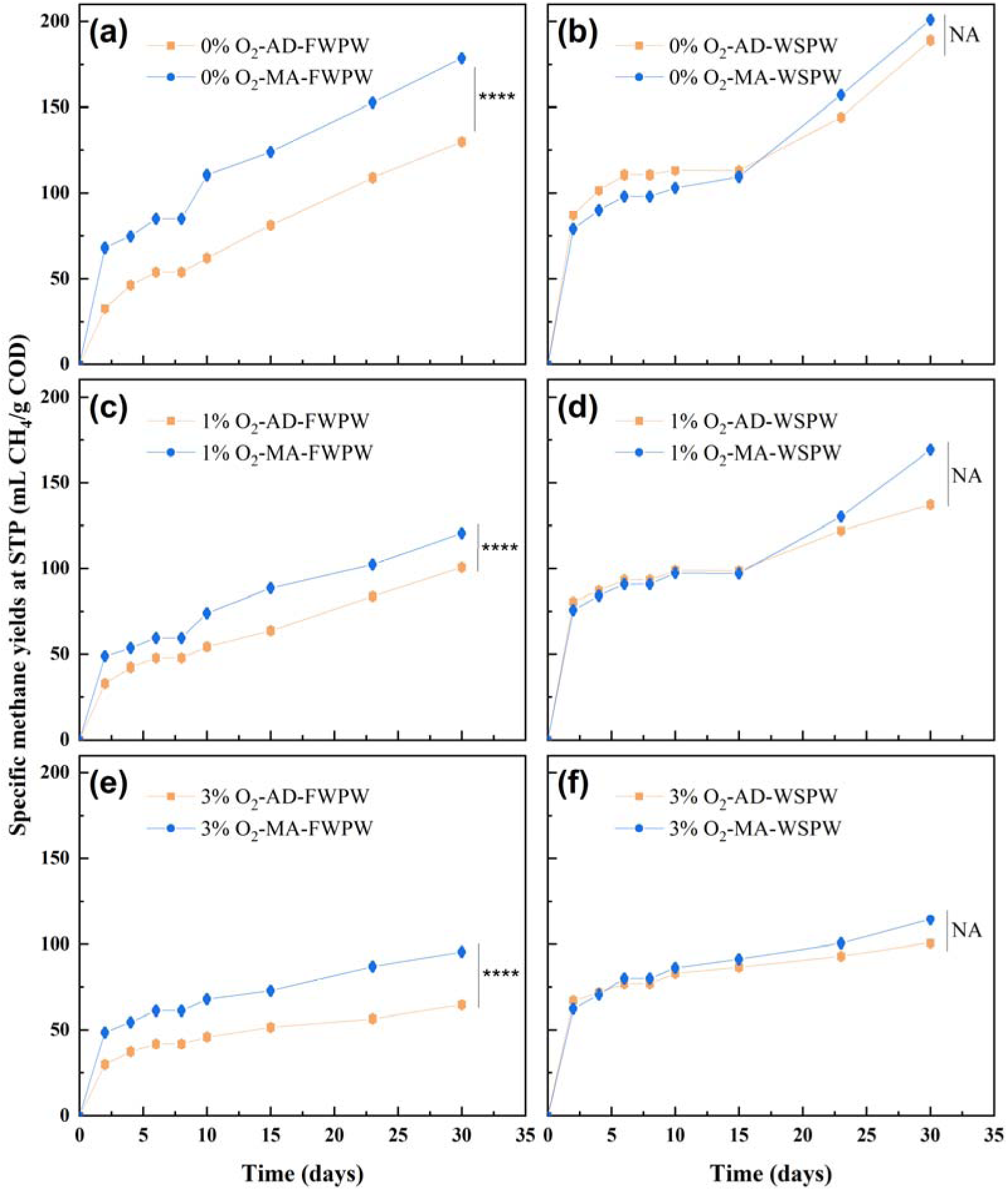
Biochemical methane potential (BMP) assays for the effects of anaerobic digestion (AD) and microaeration-acclimated (MA) biomass on biodegradation of food-waste process water (FWPW) (**a, c, e**) and wheat-straw process water (WSPW) (**b, d, f**) at the same concentration of 1 g COD/L-liquid and varying oxygen partial pressures in the headspace. Data for each experimental condition are based on duplicate experiments using separate bottles. Statistical analysis was performed using Two-Way ANOVA, and the results are presented as mean ± standard error (data points + error bars) (p-value with asterisks in the figure, **** indicates < 0.0001, NA indicates no significant difference). Some error bars may not be fully visible due to overlapping data points; detailed numerical ranges can be found in Table S7.

*In-situ* microaeration may have directly enriched facultative bacteria by promoting syntrophic interactions among the microbial groups [13, 15, 37]. However, the relationship between these microbial community shifts and the degradation of HTL process water has been poorly explored. Si *et al* [10] pointed out that the degradation of N-heteroaromatic compounds in their anaerobic reactors was likely due to oxygen introduction during feeding at the early setup stage. They observed the presence of *Pseudomonas* sp. and *Bacillus* sp., which are aerobes or facultative anaerobes capable of degrading N-heteroaromatic compounds, but did not detect anaerobes such as *Eubacterium hallii* or *Azoarcus* sp.. *Pseudomonas* sp. has also been reported to biodegrade oleate into methane under aerobic and microaerophilic conditions but not anaerobic conditions [38]. An in-depth analysis is still necessary to understand the mechanism of *in-situ* microaeration for HTL process-water degradation.

## 4 Conclusions

This study comprehensively analyzed the AD toxicity and biodegradability of HTL process water from two typical feedstocks. HTL process water inhibited methanogenesis but allowed complete glucose degradation and effective production of lactate and SCCAs during acidogenesis. Food-waste process water did not considerably inhibit the acetoclastic methanogenic activity, whereas wheat-straw process water exhibited toxicity. The toxicity of wheat-straw process water was attributed to its higher concentration of aromatic compounds without N in the rings rather than N-heteroaromatic compounds. Despite the toxicity, wheat-straw process water produced higher methane yields than food-waste process water at COD loads below 2 g/L. These findings highlight that enhancing the biodegradability of HTL process water from protein-rich feedstocks is more critical than mitigating toxicity.

In addition, we demonstrated that the feasibility of applying microaeration to enhance the AD of HTL process water is highly dependent on the characteristics of HTL feedstock. *Prior* biomass acclimation to microaeration did not increase the tolerance of acetoclastic methanogens to the toxicity of food-waste process water, and it even reduced their tolerance to wheat-straw process water. However, microaeration-acclimated biomass improved the biodegradation of food-waste process water and increased methane yields, particularly in the absence of O during the BMP assay. However, no improvement in methane yields was observed for wheat-straw process water with microaeration-acclimated biomass. These findings show that AD processes should be optimized for each different type of HTL process water.

## Author contributions

Mei Zhou: Conceptualization, Investigation, Visualization, Writing - Original Draft. Joseph G. Usack: Supervision, Conceptualization, Writing - Review & Editing, Funding acquisition. Aidan Mark Smith: Resources, Writing - Review & Editing. Largus T. Angenent: Supervision, Conceptualization, Project Administration, Writing - Review & Editing, Funding acquisition.

## Declaration of AI-assisted technologies in scientific writing

The first author used AI-assisted technologies to revise the initial manuscript draft to provide clearer and more concise English expressions. After using this tool, the first author reviewed and edited the content again and took full responsibility for the content of the published article.

## Funding and acknowledgments

This work was supported, in part, by a grant from the German-Israeli Foundation for Scientific Research and Development (GIF) (grant number I-1547-500.15/2021). L.T.A. was supported, in part, by the Alexander von Humboldt Foundation in the framework of the Alexander von Humboldt Professorship. M.Z. would like to acknowledge the China Scholarship Council (CSC, No. 202006310033) for its support. A.M.S. was supported by the Poul Due Jensen Foundation. The authors would like to acknowledge Dr. Libera Lo Presti for her support in editing.

