## Supporting Information for "Toxicity and Biodegradation of Two Different Hydrothermal Liquefaction Process Waters to Anaerobic Digestion and the Effect of Microaeration"

### Supplementary information

**Table S1** Characteristics of process water solutions and biomass.

| Materials | Food-waste process water | Wheat-straw process water | Anaerobic Inoculum | Anaerobic Biomass | Microaerobic Biomass |
|---|---|---|---|---|---|
| Color | Orange (tea-like) | black | - | - | - |
| Chemical Oxygen Demand (COD) (g/L) | 41.8 ± 0.57 | 70.1 ± 2.43 | 91.0 | 38.5 | 36.9 |
| pH | 5.23 | 5.08 | 7.8 | 7.0 | 7.0 |

**Table S2** Experimental setup of the acidogenic AT assay.

| ID | Blank | Control | FWPW_10% | FWPW_30% | FWPW_50% | FWPW_70% | FWPW_100% | WSPW_10% | WSPW_30% | WSPW_50% | WSPW_70% | WSPW_100% |
|---|---|---|---|---|---|---|---|---|---|---|---|---|
| Inoculum (mL) | 5.99 | 5.99 | 5.99 | 5.99 | 5.99 | 5.99 | 5.99 | 5.99 | 5.99 | 5.99 | 5.99 | 5.99 |
| Glucose Solution (mL) | 0 | 2.41 | 2.41 | 2.41 | 2.41 | 2.41 | 2.41 | 2.41 | 2.41 | 2.41 | 2.41 | 2.41 |
| Process Water (mL) | 0 | 0 | 0.36 | 1.08 | 1.79 | 2.51 | 3.59 | 0.21 | 0.64 | 1.07 | 1.50 | 2.14 |
| Deionized Water (mL) | 144.0 | 141.6 | 141.2 | 140.5 | 139.8 | 139.1 | 138.0 | 141.4 | 141.1 | 140.5 | 140.1 | 139.5 |
| Total Volume (mL) | 150 | 150 | 150 | 150 | 150 | 150 | 150 | 150 | 150 | 150 | 150 | 150 |
| Increased COD from process water (g/L) | 0 | 0 | 0.1 | 0.3 | 0.5 | 0.7 | 1 | 0.1 | 0.3 | 0.5 | 0.7 | 1 |
| Total COD of substrates (g/L) | 0 | 1 | 1.1 | 1.3 | 1.5 | 1.7 | 2 | 1.1 | 1.3 | 1.5 | 1.7 | 2 |

**Table S3** Experimental setup of the methanogenic AT assay.

| ID | Blank | Control | FWPW_10% | FWPW_30% | FWPW_50% | FWPW_70% | FWPW_100% | WSPW_10% | WSPW_30% | WSPW_50% | WSPW_70% | WSPW_100% |
|---|---|---|---|---|---|---|---|---|---|---|---|---|
| Inoculum (mL) | 9.9 | 9.9 | 9.9 | 9.9 | 9.9 | 9.9 | 9.9 | 9.9 | 9.9 | 9.9 | 9.9 | 9.9 |
| The mixture of Acetate and propionic acid (mL) | 0 | 30.9 | 30.9 | 30.9 | 30.9 | 30.9 | 30.9 | 30.9 | 30.9 | 30.9 | 30.9 | 30.9 |
| Process Water (mL) | 0 | 0 | 1.08 | 3.23 | 5.38 | 7.53 | 10.76 | 0.64 | 1.93 | 3.21 | 4.49 | 6.42 |
| Deionized Water (mL) | 140.1 | 109.1 | 108.1 | 105.9 | 103.8 | 101.6 | 98.4 | 108.5 | 107.2 | 105.9 | 104.7 | 102.7 |
| Total Volume (mL) | 150 | 150 | 150 | 150 | 150 | 150 | 150 | 150 | 150 | 150 | 150 | 150 |
| Increased COD from process water (g/L) | 0 | 0 | 0.3 | 0.9 | 1.5 | 2.1 | 3 | 0.3 | 0.9 | 1.5 | 2.1 | 3 |
| Total COD of substrates (g/L) | 0 | 3 | 3.3 | 3.9 | 4.5 | 5.1 | 6 | 3.3 | 3.9 | 4.5 | 5.1 | 6 |

**Table S4** Experimental setup of biochemical methane potential (BMP) assays.

| | Anaerobic biomass | | | | | | | Microaeration-acclimated biomass | | | | | | |
|---|---|---|---|---|---|---|---|---|---|---|---|---|---|---|
| ID | Blank | 0%-AD-FWPW | 1%-AD-FWPW | 3%-AD-FWPW | 0%-AD-WSPW | 1%-AD-WSPW | 3%-AD-WSPW | Blank | 0%-MA-FWPW | 1%-MA-FWPW | 3%-MA-FWPW | 0%-MA-WSPW | 1%-MA-WSPW | 3%-MA-WSPW |
| Inoculum (mL) | 2.73 | 2.73 | 2.73 | 2.73 | 2.73 | 2.73 | 2.73 | 2.85 | 2.85 | 2.85 | 2.85 | 2.85 | 2.85 | 2.85 |
| Process Water (mL) | 0 | 0.72 | 0.72 | 0.72 | 0.43 | 0.43 | 0.43 | 0 | 0.72 | 0.72 | 0.72 | 0.43 | 0.43 | 0.43 |
| Mineral Solution (mL) | 0.3 | 0.3 | 0.3 | 0.3 | 0.3 | 0.3 | 0.3 | 0.3 | 0.3 | 0.3 | 0.3 | 0.3 | 0.3 | 0.3 |
| Trace Element Solution (mL) | 0.3 | 0.3 | 0.3 | 0.3 | 0.3 | 0.3 | 0.3 | 0.3 | 0.3 | 0.3 | 0.3 | 0.3 | 0.3 | 0.3 |
| $Na_2S$ Solution (mL) | 0.3 | 0.3 | 0.3 | 0.3 | 0.3 | 0.3 | 0.3 | 0.3 | 0.3 | 0.3 | 0.3 | 0.3 | 0.3 | 0.3 |
| Anaerobic Water (mL) | 26.37 | 25.65 | 25.65 | 25.65 | 25.94 | 25.94 | 25.94 | 26.25 | 25.53 | 25.53 | 25.53 | 25.82 | 25.82 | 25.82 |
| Air in the Headspace (mL) | 0 | 0 | 1.43 | 4.28 | 0 | 1.43 | 4.28 | 0 | 0 | 1.43 | 4.28 | 0 | 1.43 | 4.28 |

**Table S5** The numerical range of error bars in Fig. 2.

| Days | FWPW_10% | FWPW_30% | FWPW_50% | FWPW_70% | FWPW_100% | Days | WSPW_10% | WSPW_30% | WSPW_50% | WSPW_70% | WSPW_100% |
|---|---|---|---|---|---|---|---|---|---|---|---|
| | Fig. 2a | | | | | | Fig. 2d | | | | |
| 1 | 0.007 | 0.003 | 0.004 | 0.001 | 0.002 | 1 | 0.002 | 0.019 | 0.007 | 0.036 | 0.037 |
| 2 | 0.007 | 0.096 | 0.004 | 0.122 | 0.002 | 2 | 0.221 | 0.108 | 0.047 | 0.232 | 0.085 |
| 3 | 0.140 | 0.027 | 0.058 | 0.358 | 0.034 | 3 | 0.473 | 0.590 | 0.215 | 0.287 | 0.096 |
| 4 | 0.504 | 0.572 | 0.330 | 0.594 | 0.233 | 4 | 1.499 | 1.211 | 0.294 | 0.302 | 1.113 |
| 5 | 0.569 | 0.237 | 0.105 | 0.890 | 0.139 | 5 | 5.517 | 2.067 | 1.172 | 2.051 | 1.938 |
| 6 | 0.556 | 0.011 | 0.750 | 1.157 | 0.229 | 6 | 2.717 | 1.768 | 0.605 | 2.321 | 1.669 |
| 7 | 0.163 | 0.062 | 0.133 | 0.963 | 0.275 | 7 | 2.371 | 0.911 | 0.934 | 2.610 | 1.725 |
| 8 | 0.536 | 0.310 | 0.108 | 0.215 | 0.572 | 8 | 2.798 | 1.090 | 1.024 | 0.213 | 1.591 |
| 9 | 1.027 | 0.026 | 0.074 | 0.386 | 0.070 | 9 | 2.404 | 1.032 | 0.959 | 0.407 | 0.528 |
| 10 | 1.177 | 1.067 | 0.099 | 0.116 | 0.881 | 10 | 2.432 | 0.901 | 1.001 | 0.594 | 0.322 |
| 11 | 0.606 | 1.866 | 0.324 | 0.914 | 0.949 | 11 | 2.432 | 0.901 | 1.001 | 0.594 | 0.762 |
| 12 | 0.606 | 1.866 | 0.324 | 0.914 | 0.949 | 12 | 2.432 | 0.901 | 1.001 | 0.594 | 0.823 |
| | Fig. 2b | | | | | | Fig. 2e | | | | |
| 1 | 0.002 | 0.001 | 0.001 | 0.000 | 0.000 | 1 | 0.001 | 0.006 | 0.002 | 0.008 | 0.007 |
| 2 | 0.002 | 0.029 | 0.001 | 0.027 | 0.000 | 2 | 0.074 | 0.033 | 0.012 | 0.051 | 0.017 |
| 3 | 0.047 | 0.008 | 0.015 | 0.080 | 0.007 | 3 | 0.158 | 0.179 | 0.055 | 0.064 | 0.019 |
| 4 | 0.168 | 0.173 | 0.085 | 0.132 | 0.046 | 4 | 0.500 | 0.367 | 0.075 | 0.067 | 0.218 |

| | | | | | | | | | | | |
|---|---|---|---|---|---|---|---|---|---|---|---|
| 5 | 0.190 | 0.072 | 0.027 | 0.198 | 0.027 | 5 | 1.839 | 0.626 | 0.301 | 0.456 | 0.380 |
| 6 | 0.185 | 0.003 | 0.192 | 0.257 | 0.045 | 6 | 0.906 | 0.536 | 0.155 | 0.516 | 0.327 |
| 7 | 0.054 | 0.019 | 0.034 | 0.214 | 0.054 | 7 | 0.790 | 0.276 | 0.240 | 0.580 | 0.338 |
| 8 | 0.179 | 0.094 | 0.028 | 0.048 | 0.112 | 8 | 0.933 | 0.330 | 0.262 | 0.047 | 0.312 |
| 9 | 0.342 | 0.008 | 0.019 | 0.086 | 0.014 | 9 | 0.801 | 0.313 | 0.246 | 0.090 | 0.103 |
| 10 | 0.392 | 0.323 | 0.025 | 0.026 | 0.173 | 10 | 0.811 | 0.273 | 0.257 | 0.132 | 0.063 |
| 11 | 0.202 | 0.565 | 0.083 | 0.203 | 0.186 | 11 | 0.811 | 0.273 | 0.257 | 0.132 | 0.149 |
| 12 | 0.202 | 0.565 | 0.083 | 0.203 | 0.186 | 12 | 0.811 | 0.273 | 0.257 | 0.132 | 0.161 |
| | Fig. 2c | | | | | | Fig. 2f | | | | |
| 1 | 0.012 | 0.010 | 0.010 | 0.004 | 0.006 | 1 | 0.002 | 0.019 | 0.007 | 0.036 | 0.037 |
| 2 | 0.000 | 0.154 | 0.000 | 0.229 | 0.063 | 2 | 0.222 | 0.096 | 0.046 | 0.195 | 0.053 |
| 3 | 0.220 | 0.552 | 0.616 | 0.546 | 0.151 | 3 | 0.584 | 0.485 | 0.176 | 0.108 | 0.114 |
| 4 | 1.215 | 0.517 | 0.324 | 0.326 | 0.280 | 4 | 1.208 | 0.661 | 0.269 | 0.480 | 1.044 |
| 5 | 1.456 | 0.820 | 0.474 | 1.433 | 0.289 | 5 | 4.109 | 0.913 | 1.008 | 2.085 | 2.025 |
| 6 | 0.248 | 0.227 | 0.890 | 1.187 | 0.662 | 6 | 2.995 | 0.774 | 0.835 | 0.410 | 1.917 |
| 7 | 1.965 | 1.407 | 1.458 | 0.176 | 1.407 | 7 | 0.404 | 1.330 | 1.128 | 2.329 | 0.931 |
| 8 | 0.557 | 0.370 | 1.357 | 2.670 | 0.493 | 8 | 0.461 | 0.517 | 0.093 | 2.817 | 1.550 |
| 9 | 0.363 | 0.307 | 0.200 | 0.921 | 0.804 | 9 | 0.395 | 0.078 | 0.076 | 0.201 | 1.547 |
| 10 | 0.117 | 0.852 | 0.579 | 0.318 | 0.613 | 10 | 0.061 | 0.298 | 0.091 | 0.194 | 0.441 |
| 11 | 0.212 | 0.793 | 0.430 | 1.610 | 0.112 | 11 | 0.212 | 0.793 | 0.430 | 1.610 | 0.441 |
| 12 | 0.606 | 1.866 | 0.324 | 0.914 | 0.949 | 12 | 2.432 | 0.901 | 1.001 | 0.594 | 0.823 |

**Table S6** The numerical range of error bars in Fig. 4.

| Days | FWPW | WSPW | FWPW | WSPW |
|---|---|---|---|---|
| | Fig. 4a | | Fig. 4a | |
| 2 | 0.040 | 0.012 | 0.940 | 0.129 |
| 4 | 0.219 | 0.026 | 0.110 | 0.774 |
| 6 | 0.229 | 0.039 | 0.339 | 2.092 |
| 8 | 0.229 | 0.039 | - | - |
| 10 | 0.219 | 0.027 | 1.124 | 3.831 |
| 15 | 0.296 | 0.027 | 0.475 | - |
| 23 | 0.244 | 0.578 | 0.286 | 4.865 |
| 30 | 0.131 | 1.518 | 3.338 | 10.752 |

**Table S7** The numerical range of error bars in Fig. 6.

| Days | 0% $O_2$-AD-FWPW | 0% $O_2$-MA-FWPW | 1% $O_2$-AD-FWPW | 1% $O_2$-MA-FWPW | 3% $O_2$-AD-FWPW | 3% $O_2$-MA-FWPW | 0% $O_2$-AD-WSPW | 0% $O_2$-MA-WSPW | 1% $O_2$-AD-WSPW | 1% $O_2$-MA-WSPW | 3% $O_2$-AD-WSPW | 3% $O_2$-MA-WSPW |
|---|---|---|---|---|---|---|---|---|---|---|---|---|
| | Fig. 6a | | Fig. 6b | | Fig. 6c | | Fig. 6d | | Fig. 6e | | Fig. 6f | |
| 2 | 0.040 | 0.117 | 0.354 | 0.046 | 0.053 | 0.259 | 0.012 | 0.498 | 0.164 | 0.112 | 0.230 | 0.108 |
| 4 | 0.219 | 0.118 | 0.374 | 0.042 | 0.077 | 0.213 | 0.026 | 0.526 | 0.125 | 0.060 | 0.200 | 0.095 |
| 6 | 0.229 | 0.161 | 0.347 | 0.044 | 0.090 | 0.164 | 0.039 | 0.540 | 0.140 | 0.072 | 0.179 | 0.074 |
| 8 | 0.229 | 0.161 | 0.347 | 0.044 | 0.090 | 0.164 | 0.039 | 0.540 | 0.140 | 0.072 | 0.179 | 0.074 |
| 10 | 0.219 | 0.170 | 0.349 | 0.124 | 0.088 | 0.179 | 0.027 | 0.539 | 0.136 | 0.099 | 0.149 | 0.083 |
| 15 | 0.296 | 0.163 | 0.390 | 0.101 | 0.076 | 0.134 | 0.027 | 0.388 | 0.136 | 0.099 | 0.146 | 0.093 |
| 23 | 0.244 | 0.237 | 0.383 | 0.201 | 0.208 | 0.131 | 0.027 | 0.030 | 0.043 | 0.077 | 0.137 | 0.031 |
| 30 | 0.131 | 0.241 | 0.348 | 0.072 | 0.233 | 0.237 | 0.027 | 0.137 | 0.190 | 0.083 | 0.017 | 0.001 |
